# The SH Protein of Mumps Virus is a Druggable Pentameric Viroporin

**DOI:** 10.1101/2024.08.09.607002

**Authors:** Kira Devantier, Trine L. Toft-Bertelsen, Andreas Prestel, Viktoria M. S. Kjær, Cagla Sahin, Marco Giulini, Stavroula Louka, Katja Spiess, Asmita Manandhar, Katrine Qvortup, Trond Ulven, Bo H. Bentzen, Alexandre MJJ Bonvin, Nanna MacAulay, Birthe B. Kragelund, Mette M. Rosenkilde

**Affiliations:** Laboratory for Molecular and Translational Pharmacology, Department of Biomedical Sciences, University of Copenhagen, Copenhagen, Denmark; Structural Biology and NMR Laboratory (SBiNLab), The Linderstrøm-Lang Centre for Protein Science and Integrative Structural Biology at University of Copenhagen (ISBUC), Department of Biology, University of Copenhagen, Copenhagen, Denmark; Department of Neuroscience, Faculty of Health and Medical Sciences, University of Copenhagen, Copenhagen, Denmark; Department of Microbiology, Tumor and Cell Biology, Karolinska Institutet, Stockholm, Sweden; Department of Biology, University of Copenhagen, Denmark; Department of Chemistry, Faculty of Science, Computational Structural Biology Group, Bijvoet Centre for Biomolecular Research, Utrecht 3584 CS, The Netherlands; Department of Chemistry, Technical University of Denmark, DK-2800 Lyngby, Denmark; Virus and Microbiological Special Diagnostics, Statens Serum Institut, Copenhagen, Denmark; Department of Drug Design and Pharmacology, University of Copenhagen, Copenhagen, Denmark; Department of Biomedical Sciences, University of Copenhagen, Denmark

**Keywords:** Porin, chloride channel, NMR, BIT225, membrane protein, HADDOCK, membrane transport, targeting, infection

## Abstract

Viral infections are on the rise and drugs targeting viral proteins are needed. Viroporins constitute a growing group of virus-encoded transmembrane oligomeric proteins that allow passage of small molecules across the membrane. Despite sparsity in viroporin structures, recent work has revealed diversity in both the number of transmembrane helices and oligomeric states. Here we provide evidence that the small hydrophobic protein (SH) from mumps virus is a pentameric viroporin. From extensive biophysical data, a HADDOCK model of full-length SH shows its intracellular C-terminal region to form an extended structure crucial to stabilization of the pentamer. Heterologous expression of wild type SH and variants in *Xenopus laevis* oocytes reveals the viroporin as a chloride channel, facilitated by conserved hydroxyl-carrying residues lining the pore. The channel function of SH is inhibited by the small-molecule BIT225, highlighting the potential for antiviral targeting through SH.

## Introduction

Mumps virus (MuV) is a highly infectious virus known to cause inflammatory symptoms, commonly parotitis and orchitis (1). MuV is neurotropic and can cause central nervous system infections or long-term effects such as seizures, deafness, or infertility (1, 2). MuV belongs to the *Paramyxoviridae* family and is an enveloped, negative-sense RNA virus (1). It is one of three viruses targeted by the MMR (measles, mumps, and rubella) vaccine; a historically highly efficient vaccine used in vaccination programs since the 1960s (1). Despite this, recent sporadic outbreaks have been observed, even in majorly vaccinated populations (2–4), reviving a focus on MuV. To combat these outbreaks, new options for treatment after infection, as a complement to vaccination, could be considered.

The small hydrophobic protein (SH) is one of the nine proteins encoded by MuV. SH is a 57-residue single-pass type I membrane protein, with the C-terminus residing in the cytosol (5) (Fig. 1a-b). It is not fully known which membranes SH locates to, with some suggesting that SH is only found in infected host cells, such as the plasma membrane (5), and not on the surface of circulating MuV (1). The gene encoding SH is in one of the more variable parts of the MuV genome, and its sequence is used to distinguish between the twelve genotypes of MuV (6). In recent years, the most prevalent genotype for recorded MuV cases has been genotype G (4).

**Figure 1.**
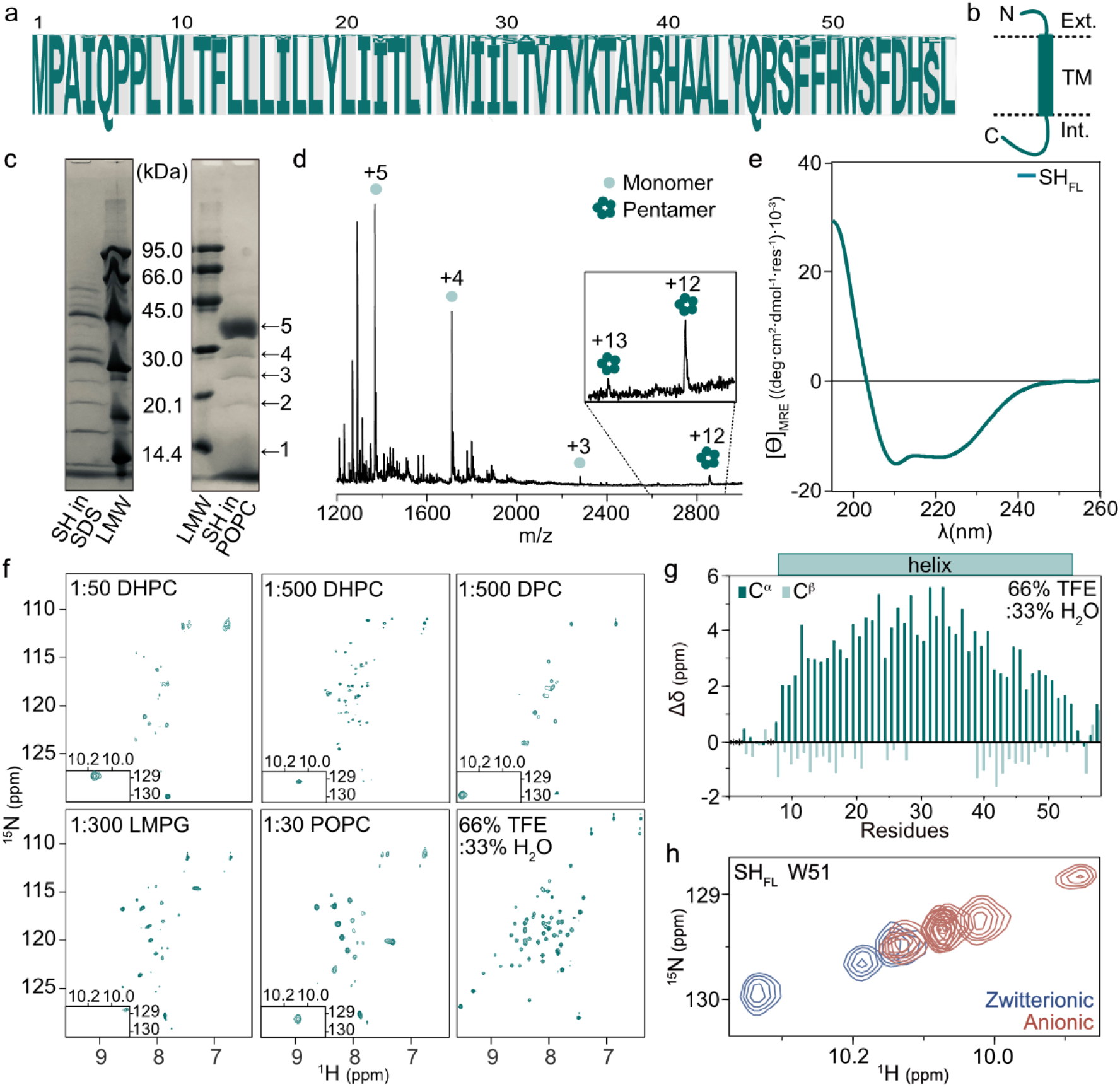
Oligomerization and structural characterization of SH_FL_. a) Consensus amino acid sequence of SH protein from genotype G. b) Membrane topology of SH. c) Left: SDS-PAGE of SH_FL_ in 100 mM SDS. Right: SDS-PAGE of SH_FL_ in 1:120 POPC. d) Native mass spectrometry of SH_FL_ in DPC. e) Far-UV CD spectrum of SH_FL_ in 1:500 DHPC. f) ^1^H-^15^N HSQC spectra of SH_FL_ in different membrane mimetics and detergents as indicated. g) SCS analysis of SH_FL_ in 66% TFE:33% H_2_O (v/v). h) ^1^H-^15^N HSQC spectra of the Trp51 indole peak from (*f*). Peaks coloured by lipid or detergent head group charge. Anionic (red); Zwitterionic (blue).

The overall effects of SH in the viral life cycle in humans, its single natural host, are not well understood. Studies in cell cultures have shown that SH is not essential for viral replication (5), but can contribute to viral pathogenesis (7). MuV SH have been shown to interfere with TNF-α-mediated apoptosis (8), and be interchangeable with SH from related *paramyxoviruses* human respiratory syncytial virus (hRSV), simian virus 5, and J paramyxovirus (JPV), despite low sequence similarity (9, 10). Of interest, hRSV SH has been described as a viroporin (11, 12). Viroporins are a family of virus-encoded membrane proteins that form pores in virions and host membranes, altering the permeability to certain ions, leading to successful propagation of viral infections (13, 14). Only a limited number of viroporins have been extensively characterised. Notable examples of these are M2 of influenza A virus (15, 16) and E protein of SARS-CoV-2 (17, 18). M2 and E display a diversity of structural states, both within the individual viroporin and compared to each other and other viroporin structures (14). Viroporins are known to have a diverse set of functions with actions in multiple steps of the viral life cycle, including entry to egress as well as immune evasion, and they are likely underutilised as targets for antiviral drugs (14, 19).

Given the resemblance of MuV SH to viroporins, we address here whether SH has viroporin functions. We use an interdisciplinary approach to reveal SH as an oligomeric chloride channel and provide an integrative structural model of the pentameric pore formed by full-length SH. Using mutagenesis, we identify key determinants for the channel function of SH as well as for stabilizing its pentameric state. In demonstrating the possible inhibition of SH by a small molecule inhibitor, the potential use of SH as a drug target in antiviral defence is revealed. Collectively, our data establish SH as a viroporin.

## Results

### SH forms oligomers

Many membrane proteins are known to oligomerize, including viroporins and the related viroporin hRSV SH (11). We therefore assessed the oligomeric state of SH using several approaches. First, we recombinantly produced full-length SH (SH_FL_) allowing stable isotope enrichment for nuclear magnetic resonance (NMR) spectroscopy analyses. Analysing pure SH_FL_ on SDS-PAGE, showed several bands distributed as a ladder with step sizes corresponding to monomeric SH_FL_ of an expected size of 6.8 kDa (Fig. 1c, left). This finding indicated that SH_FL_ forms oligomers, with the largest size corresponding to that of a decamer. To investigate which oligomeric state(s) of SH_FL_ were dominant and even stabilized in lipids, SH_FL_ reconstituted in 1:120 1-palmitoyl-2-oleoyl-glycero-3-phosphocholine (POPC) was investigated by SDS-PAGE, increasing the sample buffer SDS from 2% to 4% (w/v) and refraining from sample heating (Fig. 1c, right). Here, we observed that SH_FL_ primarily migrated to the apparent size of ∼34 kDa, corresponding to a pentamer. From the pattern of consecutive peaks, native mass spectrometry (MS) of SH_FL_ in detergent also indicated a pentameric state (Fig. 1d). Thus, the dominant oligomeric state of SH_FL_ in multiple environments is that of a pentamer.

### SH_FL_ is helical in reconstituted environments

To determine the structure of SH_FL_, far-UV circular dichroism (CD) spectra of SH_FL_ in 1,2-dicaproyl-sn-glycero-3-phosphocholine (DHPC) at a 1:500 protein:lipid ratio were recorded at 25° C. The spectra showed a characteristic signature of α-helical structure of the SH_FL_ (Fig. 1e). The fractional helicity (f_H_) based on the mean residue ellipticity at 222 nm (MRE_222_) was 43% corresponding to 25 residues. In contrast, AlphaFold3 predictions of SH predict the protein to be mostly helical, ≥48 residues/84%, in both monomeric and pentameric models (Fig. S1), although with low local distance difference test (pLDDT) scores and an interface predicted template modelling (ipTM) of 0.26 for the pentameric model. SH_FL_ was tested in a variety of membrane mimetics and displayed helical properties in all conditions with f_H_ ∼40% (Fig. S2a). To obtain a higher level of structural information for SH, a detergent and lipid screen with commonly used detergents and lipids was performed by NMR using ^15^N-labeled SH_FL_. For all detergents and lipids, the corresponding ^1^H-^15^N HSQC spectra of SH_FL_ showed only few peaks compared to the expected number for SH_FL_ (Fig. 1f) suggesting a higher-order structure. The two spectra of SH_FL_ reconstituted in lipids, 1-myristoyl-2-hydroxy-sn-glycero-3-phospho-(1′-rac-glycerol) (LMPG) and POPC, had slightly more peaks compared to those in detergents; however, none of these conditions gave spectra that would allow assignment and further structural characterization. Close to the expected number of peaks were observed for SH_FL_ in 66% trifluoroethanol (TFE):33% H_2_O (v/v) and assignments of the backbone and sidechain nuclei of SH_FL_ were possible using a ^13^C,^15^N-labeled sample (Fig. S3). Secondary chemical shift (SCS) analysis, comparing the obtained chemical shifts to those of predicted random coil of SH_FL_, revealed that SH_FL_ was helical throughout most of the sequence, including residues 8-54, leaving only the N- and C-terminal 1-7 and 55-57 residues disordered (Fig. 1g). As TFE is known to induce helicity (20), the effect of varying concentrations of TFE was explored. A correlation between increasing TFE concentration and fractional helicity in SH was observed (Fig. S2b), suggesting that the helicity of SH in high TFE was artificially increased compared to the native protein structure. One of the few peaks visible in every spectrum was from a Trp side chain indole, which showed chemical shift changes depending on the solubilizing conditions (Fig. 1h). As SH_FL_ contains two Trp residues, Trp27 and Trp51, with Trp27 in the TM region, the observable peak most likely corresponds to Trp51, indicating it is close enough to the membrane-water interface to be affected by the type of the membrane mimetics.

### The helicity of SH_FL_ is confined to the TM

To address the effect of the C-terminus, an SH variant with a truncated C-terminus (SH_1-34_), was purified and investigated by CD and NMR and compared to SH_FL_. Here, the ^1^H-^15^N HSQC spectrum of SH_1-34_ in 1:500 DHPC displayed the expected number of peaks. Native MS of SH_1-34_ revealed a change in oligomeric state to a tetrameric state, further supported by SDS-PAGE (Fig. S4a,c). Thus, the C-terminus of SH is important for stabilizing the pentamer. The spectral quality made backbone assignment possible (Fig. 2a), allowing for SCS analysis and characterization of the dynamics of SH_1-34_ (Fig. 2b). SCS analysis showed α-helical content from residue 8-31, without distinct secondary structure in either terminus. The recorded NMR relaxation experiments revealed that SH_1-34_ displays low *R*_*1*_, high *R*_*2*_ and HetNOE values in the region where the protein is helical (Fig. 2b), corresponding to the region of the transmembrane (TM) domain (TMD). By CD, the profile of SH_1-34_ was α-helical (Fig. 2c) with an f_H_ of 75% compared to 43% for SH_FL_. Converted to number of residues, both variants have 25 residues in the helical conformation. In conclusion, only the TM-part of SH is helical, whereas the C-terminal tail has non-helical structure that exerts a stabilizing effect on the pentamer and likely in proximity to the membrane.

**Figure 2.**
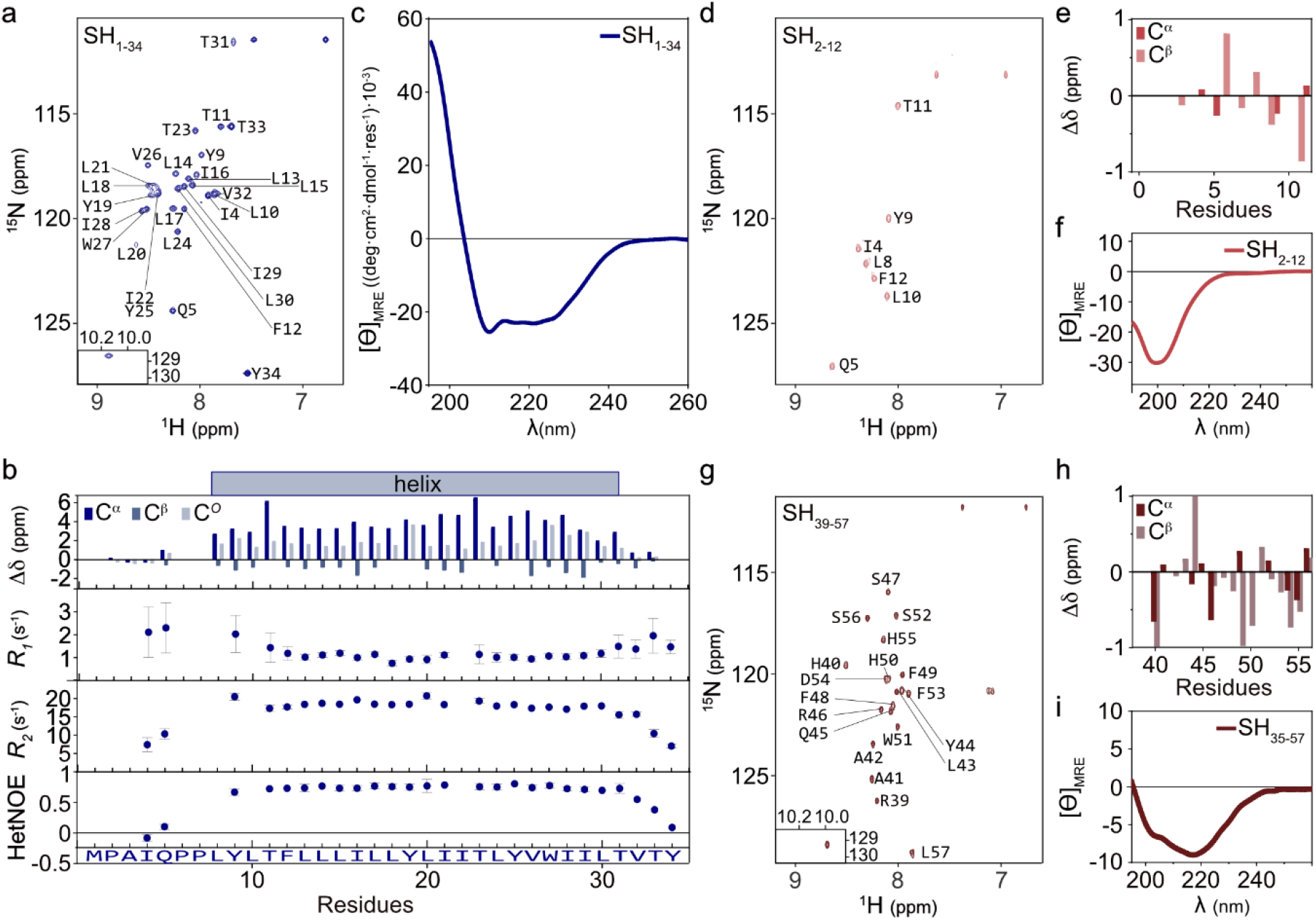
SH is helical only in its TMD. a) ^1^H-^15^N HSQC spectrum of SH_1-34_ in DHPC, with assignment. b) SCS analysis of SH_1-34_ in (*a*). *R*_*1*_, *R*_*2*_ and HetNOE values for SH_1-34_ in DHPC. c) Far-UV CD spectra of SH_1-34_ in DHPC. d) ^1^H-^15^N HSQC spectrum of SH_2-12_, with assignment. e) SCS analysis of SH_2-12_ in (*d*). f) Far-UV CD spectra of SH_2-12_. g) ^1^H-^15^N HSQC spectrum of SH_39-57_, with assignment. h) SCS analysis of SH_39-57_ in (*g*). i) Far-UV CD spectra of SH_35-57_.

### The extramembrane regions of SH_FL_ show disordered and extended structural properties

Since the comparative analyses of the helical content suggested that SH_FL_ contains non-helical regions, we sought to address how the N- and C-terminal regions would behave in isolation. An N-terminal peptide of the first 12 residues, SH_2-12_, was characterised by NMR and CD spectroscopy (Fig. 2d-f). From NMR SCS analysis (Fig. 2e) a lack of secondary structure within the SH_2-12_ N-terminus was evident, supported by a distinct minimum at 198 nm in the CD spectrum (Fig. 2f). Thus, the N-terminus of SH is likely disordered up to the beginning of the TM region around residue 8. A similar analysis was performed for C-terminal peptides, SH_35-57_ or SH_39-57_ (Fig. 2g-i). No indication of a defined secondary structure was observed for SH_39-57_ (Fig. 2h). As opposed to SH_2-12_, the CD spectrum of SH_39-57_ had a minimum at 218 nm indicating an extended structure (Fig. 2i). Neither POPC nor a combination of POPC/palmitoyl-oleoyl phosphatidylserine (POPS) altered the CD profile of SH_39-57_ suggesting that the presences of a membrane surface does not induce secondary structure (Fig. S2c). Collectively, these data show that the helicity of SH is restricted to the TM region, whereas the extra membranous regions are more disordered (N-terminal), or form extended conformations (C-terminal).

### SH forms a pore with polar residues exposed towards the lumen

Using the multibody docking capabilities of HADDOCK (21–23), we next integrated the information obtained from the above experiments to generate a structural model of SH. First, a model of SH residues 7-34 (SH_7-34_), corresponding to the TM region (Fig. 2b) was predicted with AlphaFold2 v1.5.2 (24, 25) showing pLDDT scores consistently above 70 (Fig. S5a). This model was used as a starting point for the reconstruction of the multimer form of SH. Five copies of SH_7-34_ were included in the model, imposing C5 symmetry between chains, and with center-of-mass restraints and additional torsion angles restraints derived from the NMR data for SH_1-34_. The top-ranking resulting structure clusters all generated a pore exposing most of the polar residues of SH facing inward towards the lumen of the pore, including Tyr19 and Thr23 (Fig. 3a-c and S6). SH_FL_ and SH_1-34_ were also modelled as pentamers using AlphaFold3 (Fig. S1b and S5b), however all models had pLDDT scores below 70 throughout the sequence, ipTM scores below 0.5 and did not consistently position the polar residues towards the pore (Fig. S5b), likely highlighting the sparsity of viral proteins in the PDB. We thus continued with the data-supported model of SH_7-34_. The cross-section of the HADDOCK generated pore was analysed using the HOLE program (26) (Fig. 3b-c), showing the pore to be narrower towards the N-terminal end, with Phe12 and Tyr19 creating the smallest cross-sections of the pore. The interfaces between adjacent helices are dominated by extensive hydrophobic interactions, such as the interactions of methyl-rich side chains in the region of Ile16 of helix *i* and Leu15 and Leu18 on helix *i*+1, as well as Thr23 and Leu24 of helix *i* and Val26 on helix *i*+1 (Fig. 3d). Aromatic side chains also partake in the stabilization of the helix-helix interface, e.g., through a carbon-π interaction between Phe12 from helix *i* and Thr11 *i+1*. The interface-oriented Trp27 of helix *i* to form a cross-helix hydrogen bond with the hydroxyl of Thr33 on helix *i+1*, in addition to forming hydrophobic interactions with Ile29 (Fig. 3d).

**Figure 3.**
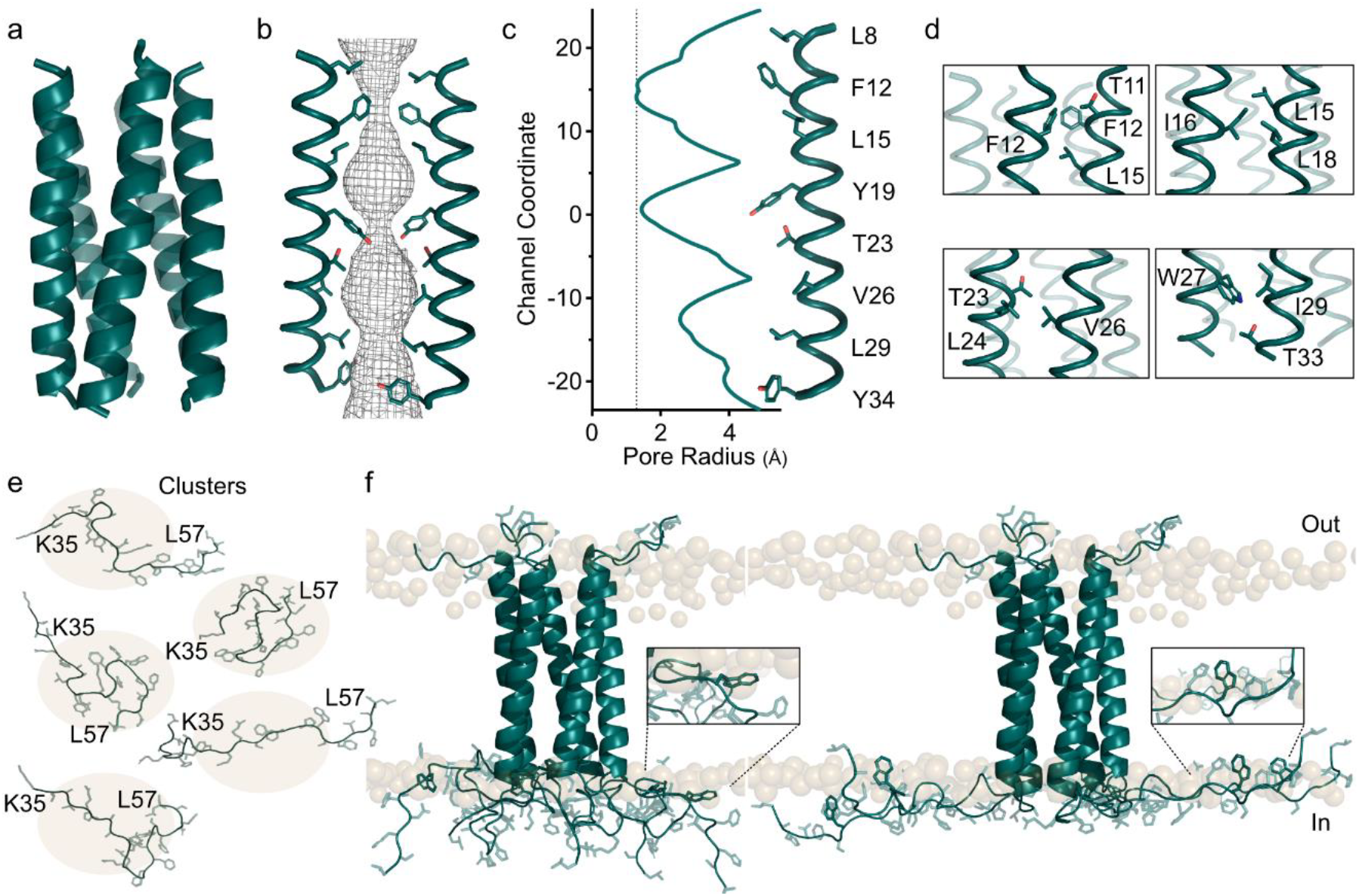
The SH pentamer. a) Five helix bundle of SH transmembrane region (7-34). b) Simplified two helix view of pore-lining residues and HOLE-calculated pore diameter. c) Pore radius of SH_7-34_ along with pore-lining residues. d) Examples of helix stabilizing residues in the helix-helix interface. e) Clusters from MD simulation of SH_35-57_. f) Two pentameric models of SH_FL_ with extended C-termini. Insert: Zoom of Trp51 proximity to “membrane”.

### The extramembrane regions of SH are extended

The N-terminal region of the SH_FL_ AlphaFold3 model (residues 1-6) displayed disorder (Fig. S1a). Disorder in the N-terminus is in full accordance with the experiments (Fig. 2b, d-f). Therefore, the six-residue N-terminal peptide was joined to residue seven of the SH_7-34_ segments using PyMoL (27) and torsion angles optimized using Coot (28) (Fig. 3f). In contrast, the C-terminal region of the SH_FL_ AlphaFold3 models (Fig. S1), SH_35-57_, was not in agreement with experimental CD and NMR data (Fig. 1e and Fig. 2g-i). The AlphaFold3 model showed continued helical structure with pLDDT scores above 70 (Fig. S1a). The experimental data clearly demonstrated a lack of helical secondary structure, and a 100 ns molecular dynamics (MD) simulation was run on the isolated SH_35-57_ peptide. Based on Root Mean Square Deviations (RMSD), conformations of the C-terminal peptide were clustered (Fig. 3e). Clusters were excluded if there was steric hindrance to residue 35, which needed to be available to form a peptide bond with residue 34 of the TM region. Five clusters of mostly extended structures, matching the observation from CD and NMR, were used for a second round of multibody docking in HADDOCK, docking five copies of SH_35-57_ to the generated TM pentamers, imposing distance restraints between residue 34 on SH_7-34_ and residue 35 on SH_35-57_ to restore the peptide connectivity. A shape consisting of fake beads representing the phosphates in the head groups of a POPC lipid bilayer was added to the molecular docking as a guideline of steric hindrance imposed by the bilayer. The resulting clusters showed diversity in the orientation of SH_35-57_, with the C-termini of some clusters colliding with the membrane, excluding these from further consideration (Fig. S7). Since the NMR spectra of SH_FL_ suggested the indole group of Trp51 to be in proximity of the membrane surface, an ambiguous restraint was placed on Trp51 of SH_35-57_ to be within 2 Å of any shape bead on the C-terminal side of the bilayer, further restricting the movement of SH_35-57_. Two different clusters fulfilled the set criteria, in which most of the C-terminal regions are extended near the membrane surface (Fig. 3f), and where Trp51 is near the membrane (Fig. 3f inset). Thus, in conclusion, our structural model of SH_FL_ indicates a transmembrane helix with a disordered N-terminus and an extended C-terminus in proximity to the membrane surface, forming pentamers, stabilized by the presence of the C-terminus.

### SH is a viroporin that conducts Cl^-^ currents

From the properties of the pentameric structure, it was relevant to explore whether SH is a viroporin with ion channel properties. SH_FL_ with a C-terminal His6-tag was heterologously expressed in *Xenopus laevis* (*l*.) oocytes, a system with low background expression of other ion channels, and characterized with conventional two-electrode voltage clamp (Fig. 4a). Uninjected oocytes served as controls to detect background current. SH_FL_-expressing oocytes showed significantly higher ion conductance than uninjected oocytes (Fig. 4b-c). The amplitude of the SH_FL_-mediated current depended on the amount of RNA microinjected into the oocytes and thus the inferred expression level of the viroporin (Fig. 4d-e). Thus, heterologously expressed SH_FL_ mediate ion conductance in a dose-dependent fashion.

**Figure 4.**
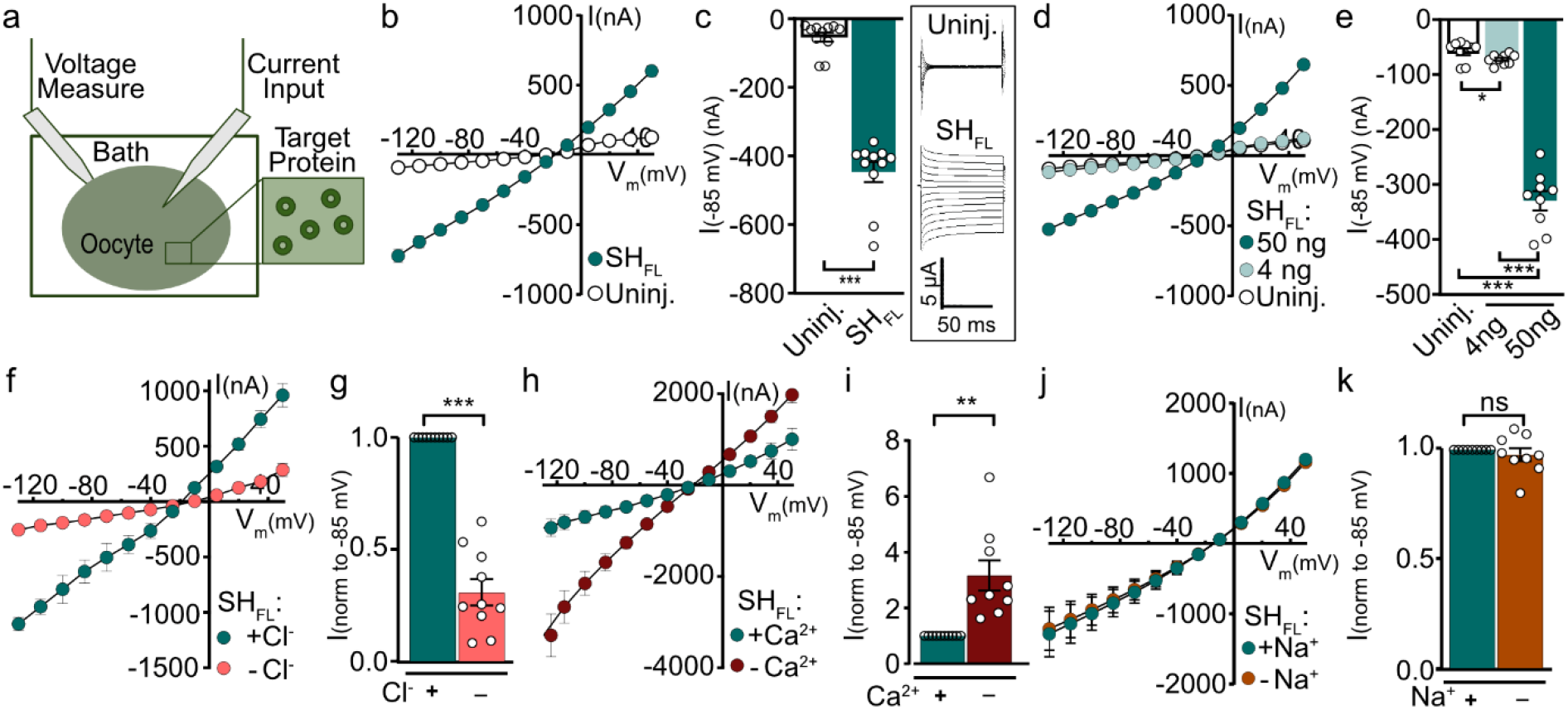
SH is a viroporin and conducts Cl^-^ currents in *Xenopus l*. oocytes. a) Illustration of the two-electrode voltage clamp setup employed. One microelectrode is used for current injection, and one is used for voltage sensing, measuring the membrane potential (V_m_) compared to a command voltage, which is levelled by an amplifier. b) Summarized and averaged current-voltage (I/V) relations in SH_FL_-expressing oocytes compared to control (uninjected) oocytes. n = 11. c) Current activity at −85 mV in SH_FL_-expressing oocytes compared to uninjected oocytes from (*b*). Inset: representative current traces. d) Summarized current-voltage (I/V) curves of uninjected or with 4, or 50 ng SH_FL_ RNA/oocyte microinjected. n = 9. e) Current activity at −85 mV in SH_FL_-expressing oocytes compared to uninjected oocytes from (*d*). f) Summarized current-voltage (I/V) curves of SH_FL_-induced currents in the presence or absence of Cl^-^. n = 10. g) Normalized data at V_m_ = -85 mV from (*f*). h) Summarized current-voltage (I/V) curves of SH_FL_-mediated currents in the presence or absence of Ca^2+^. n = 9. i) Normalized data at V_m_ = -85 mV from (*h*). j) Summarized current-voltage (I/V) curves of SH_FL_-induced currents in the presence or absence of Na^+^. n = 7. k) Normalized data at V_m_ = -85 mV from (*j*). All summarized current traces represent mean ± SEM. Each point in the bar charts indicate an individual oocyte. The magnitude of SH_FL_-mediated currents (at V_m_ = -85 mV) was compared to those of uninjected oocytes using an unpaired t-test (*c*), one-way ANOVA (*e*), or between the presence or absence of indicated ion with paired t-test (*g,i,k*). * P < 0.033; ** P < 0.002; *** P < 0.001.

To characterize the current mediated by SH_FL_, selected ions in the control recording solution were replaced. Replacement of Cl^-^ with equi-osmolar gluconate reduced the SH_FL_-mediated current (Fig. 4f; compare -632 ± 103 nA in control solution to -165 ± 26 nA with gluconate substitution, n = 10, p < 0.001, Fig. 4g). Removal of Ca^2+^ from the control solution significantly increased the SH_FL_-mediated current (Fig. 4h; compare -583 ± 116 nA in control solution to -1689 ± 186 nA without Ca^2+^, n = 9, p < 0.01, Fig. 4i), suggesting possible modulation of the pore. Replacement of Na^+^ with equi-osmolar choline left the SH_FL_-mediated current unchanged, suggesting that the current was independent of the presence of the monovalent cation (Fig. 4j; -compare 847 ± 196 nA in control solution to -800 ± 183 nA with choline substitution, n = 7, p = 0.48, Fig. 4k). Combined, these data illustrate that the major contributor to SH_FL_-mediated current in *Xenopus l*. oocytes is Cl^-^, and the viroporin is negatively modulated by Ca^2+^.

### Oligomerisation is essential for the ion channel function of SH

To resolve the identity of the SH conducting parts and validate the HADDOCK model, we constructed a set of gradually truncated SH constructs as well as an L24A, W27A (SH_LW_) and a Y19A (SH_Y_) variant, expected to affect the oligomer (Fig. 5). Four of the five variants, SH_Y_, SH_LW_, SH_13-57_ and SH_1-34_, were less conductive compared to SH_FL_ (Fig. 5b-c). Since SH_10-57_ was conducting, but SH_13-57_ was not, the disordered N-terminal prior to the beginning of the TM helix play no role in ion conductance, whereas residues 10-13 form part of the interhelical hydrophobic network stabilizing the oligomer (Fig. 3d). SDS-PAGE analysis and changes in the NMR spectra compared to SH_FL_ showed altered dynamics in the two variants SH_LW_ and SH_1-34_ indicating that these modifications affected the stability of oligomeric state, explaining the lack of ion conductance (Fig. S4). The point mutation in SH_Y_ reduced the conductance compared to SH_FL_, suggesting the tyrosine residue could be involved in ion conductance. Because Y19A would make the pore wider in an otherwise unaltered pore (Fig. 5e), it is also possible that the mutation affects the overall organisation of the pore and even lead to pore collapse. Together, these data strongly support the HADDOCK model of SH, highlighting that destabilising SH either within the TMD interfaces or removing the C-terminal tail is detrimental to its pore-conducting abilities.

**Figure 5.**
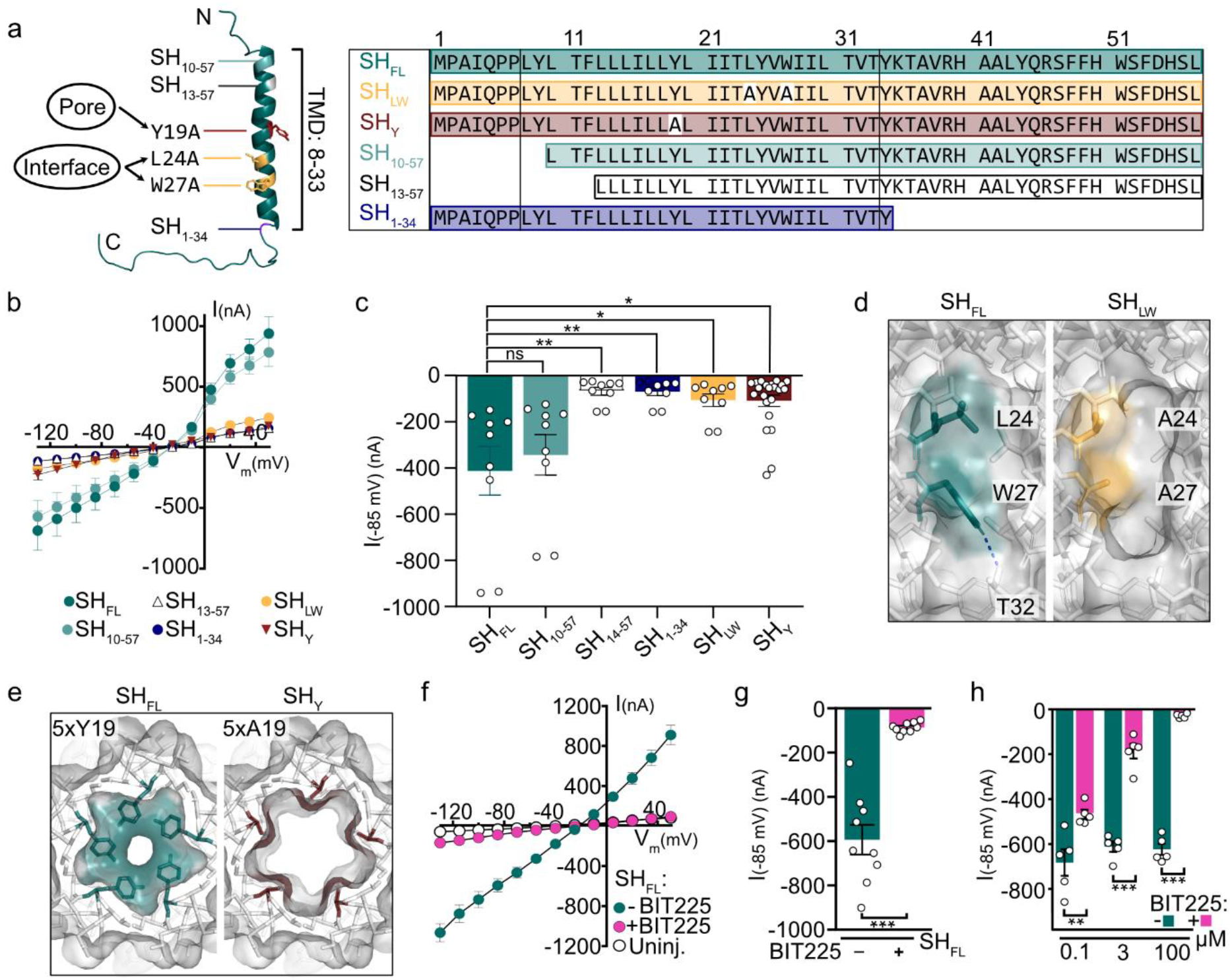
SH channel function is dependent on oligomerisation, polar residues and is inhibited by BIT225. a) Left: Schematic representation of SH variants. Right: Sequences of SH WT and variants, all electrophysiology experiments included a C-terminal His6-tag. SH_LW_: L24A, W27A. SH_Y_: Y19A. b) Summarized current-voltage (I/V) curves of SH variants. n = 9,22. c) Data at V_m_ = -85 mV from (*b*). d) Surface side-view of SH_FL_ and SH_LW_. e) Surface N-terminal-view of SH_FL_ and SH_Y_. f) Summarized current-voltage (I/V) curves of SH_FL_-expressing oocytes with or without 10 μM BIT225. n = 9. g) Data at V_m_ = -85 mV from (*f*). h) Concentration dependent inhibition of SH_FL_ by BIT225. All summarized current traces represent mean ± SEM. Each point in the bar charts indicate an individual oocyte. Statistical significance was determined with unpaired t test. * P < 0.033; ** P < 0.002; *** P < 0.001.

The amiloride analogue BIT225 is a known inhibitor of HIV-1 Vpu viroporin (29) and SARS-CoV-2 E (30) but not TMEM16, an endogenously expressed Cl^-^ ion channel in *Xenopus l*. (30). To resolve if this compound inhibits SH viroporin conductance, we preincubated SH_FL_-expressing oocytes for 45 min with 10 μM BIT225. Application of BIT225 to the oocytes resulted in a reduction in current activity (Fig. 5f-g). By varying the BIT225 concentration, it could be seen that the inhibitory effect increased with increasing concentration (Fig. 5h). Altogether these experiments showcase that SH can be targeted by small-molecule inhibitors.

## Discussion

SH has previously been identified as a type I membrane protein with implications in the immune evasion of MuV (5, 8, 9). However, like many other viral membrane proteins, structural and functional insight was lacking. Here we show that SH forms a helical pentamer with chloride conductance that is negatively modulated by calcium and inhibited by the small molecule BIT225. Through a sectional and interdisciplinary approach, a structural model of the full-length SH is provided with a dominating pentameric state observed in multiple environments (Fig. 1c-d). The resulting model shows a mostly hydrophobic TM pore, with hydrophobic side chains stabilizing the helix-helix interfaces and where a few polar residues positioned in the pore lumen, Tyr19 and Thr23, facilitate ion conduction (Fig. 5b). These data establish SH as a viroporin.

The model of SH_FL_ displays nonhelical extra-membranous regions. The six N-terminal residues show intrinsically disordered properties (31), whereas the C-terminus have extended characteristics and are tethered to the membrane surface through its many hydrophobic residues, including Trp51 (Fig. 3f). Most of the current structures of viroporins have mainly focused on the transmembrane regions of the proteins, and the few structures with extra-membranous regions generally show distinct secondary structural elements, e.g., a second α-helix on M2 from Influenza A virus perpendicular to the membrane (32). In the case of SH, no secondary structure was observed for the C-terminus and helicity could only be induced in very high concentrations of TFE (Fig. 1g and Fig. S2b). In some viroporins, short β-sheets have been suggested in the juxtamembrane regions of the protein, such as for hRSV SH (11) and SARS-CoV-2 E (18), more alike the extended structure observed for SH. Even though a lack of distinct secondary structure was observed in the C-terminus, it is expected to remain close to the membrane, as it has a high content of hydrophobic residues and displayed distinct head-group effects dependent on the lipid and detergents used (Fig. 1h). As many membrane proteins are inherently sensitive to their lipid environment(33), specific lipids might affect the conformation and activity of SH, such as affecting the organization of the C-terminus.

We identified several important features of SH. The intracellular C-terminus was directly linked to the pore function, likely through its stabilizing effect promoting the ability of SH to form oligomers. However, the exact structure and dynamics of the C-terminus is not clear. It is possible that this stabilizing effect may act through interactions between the C-terminal residues of one helix with side chains in the TM region of helices positioned (i,i±1) or (i,i±2). However, our computational modeling did not capture any of these potential contacts. As we saw no signs of intermolecular interactions between chains of the C-terminal regions, there is no evidence that these would engage in an inter chain structure rich in β-sheets, as seen for other viroporins (11, 18). A possible explanation for the stabilization of the pentamer is through entropy optimization (34). Given that the C-terminal region is extended and in contact with the bilayer and given that the C-termini do not interact with each other, optimizing the distances between them on the bilayer surface may provide an entropic force driving the pentamer together and thereby stabilizing it. Removal of the C-terminal region gave access to the structure of the TM-helix through disruption of the pentamer, into a state with favorable NMR dynamics. In contrast, removal of the N-terminus prior to the beginning of the TM-helix had no effect on conductance. Other, more specific residues were also shown to be important for the stability of the pore. The L24A and W27A mutations abolished the pore function, probably due to disruption of the network of hydrophobic residues lining the helix-helix interface, causing a similar destabilization of the pore. The Y19A mutation also showed reduced conductivity, highlighting its potential role in ion binding or in pore formation.

The oligomerisation and consequent pore function establish SH to belong to the viral protein family of viroporins. Most identified viroporins have shown a preference for cation conduction, e.g., protein 3a of SARS-CoV2 and p7 of hepatitis C virus have preferences for divalent (Ca^2+^) (35) or monovalent cations (Na^+^ and K^+^) (36), respectively. In contrast, SH displayed a preference for conducting Cl^-^ ions (Fig. 4f). This has also been observed for M2 from influenza C and D, which both contain a conserved luminal Tyr residue, where it was proposed to enhance viral budding by promoting an efflux of Cl^-^ ions, however the mechanism is unknown (37). The described effect of SH on immune evasion (8) has not yet been connected to its channel function. It is therefore possible that SH is multifunctional. The conductance of SH was negatively affected by the presence of Ca^2+^, highlighting a potential allosteric effect. It was not elucidated if and where on SH, the Ca^2+^-ions bind. A recent study on Ca^2+^ binding to disordered proteins found that binding depends on a significant amount of negatively charged residues (38). This is not a feature of either the C- or N-terminal regions of SH, although a pentameric state may jointly support Ca^2+^ binding through co-localization of up to five Asp54. As many of the lipid head groups also bind Ca^2+^ (39), one mechanism could be to destabilize the headgroup interactions of the C-terminal regions enough to weaken the pore stability and, thus, ion conductance. The effect could also be a combination of these effects, or indirectly by affecting interactions with other proteins or it may act through binding to phosphoryl groups if phosphorylation of the conserved serine residues in the C-terminal tail occurs. More data are needed to elucidate how this effect is established.

The conductivity of SH was shown to be inhibited by BIT225 (Fig. 5f), highlighting the possible druggability of SH and introducing SH as a target for anti-viral drug design. Similar druggability has been explored for viroporins from coronaviruses (40, 41). Given the importance of structure-based drug design (42), this current SH model offers insights into the organization of the oligomer and pinpoints residues important for the pore function, which could be used for specific targeting. With the recombinant production and reconstitution of the full-length SH, there is a relevant potential for initiating future high-throughput screening campaigns of larger small-molecule libraries using, e.g., saturation-transfer-difference (STD) NMR (43), which can detect selective binding of compounds at low SH concentrations.

This combined work on the structure and function of SH presents a full-length structural model of a viroporin with chloride conductance function and critical dependence on the C-terminal region for its formation, not seen for other viroporins. The data highlights SH as a potential drug target, given its implications on immune evasion and verified druggability. Besides the evidence that SH can be blocked by a generic inhibitor, we also pinpoint key residues and features in the pentameric model, including the C-terminus, that could be novel sites of interactions to be targeted by antiviral drugs.

## Acknowledgments

We thank Signe A. Sjørup, Rodrigo V. Honorato, Amer Mujezinovic and Søren Petersen for expert technical assistance. Johan G. Olsen is thanked for discussions on membrane protein complexes, and Katrine Bugge for discussion on membrane protein purification. We thank Michael Landreh for access to the Synapt G1.

## Author contributions

K.D., T.L.T-B., K.S., N.M., B.B.K., and M.M.R. conceived the study, K.D. conducted all biophysical data on SH and variants with NMR assistance from A.P., K.D analyzed all biophysical data in collaboration with B.B.K. Electrophysiological studies were conducted and analyzed by T.L.T-B., V.M.S.K. and K.S. in collaboration with N.M. and B.H.B.. C.S. conducted and analyzed native MS. K.D. conducted molecular docking with assistance from M.G. and analysis in collaboration with A.M.J.J.B.. S.L. and K.Q. synthesized SH peptide. A.M. and T.U. synthesized BIT225. K.D. designed and made all figures, with graphs from the electrophysiological studies drafted by T.L.T-B and V.M.S.K. K.D., B.B.K., and M.M.R. wrote the manuscript with input from all authors.

## Materials and methods

### Methods

#### Protein purification

Expression and purification of SH (WT and variants) for structural characterization was performed as described in Bugge et al. 2015 (44). In brief, a pGEX-4T-1 vector coding for SH downstream of a GST carrier protein and a thrombin cleavage site was acquired from GenScript (US). The plasmid was transformed into competent *E. coli* BL21(DE3) cells, and protein was expressed unlabelled in LB media. Labelled ^15^N or ^13^C,^15^N protein was expressed in M9 media (3 g/L KH_2_PO_4_, 7.5 g/L Na2HPO_4_·H_2_O, 5 g/L NaCl, 1 mM MgSO_4_, 1 mL M2 trace solution, 3 g/L glucose, 1.5 g/L (NH_4_)_2_SO_4_ (if ^13^C-labelled: ^13^C-D-Glucose (ISOTEC), if ^15^N-labelled: (^15^NH_4_)_2_SO_4_, (ISOTEC)) added 100 μg mL^-1^ ampicillin (Fisher BioReagents). Cells were collected by centrifugation (15 min, 5000*g*, 4 °C) and resuspended in 40 mL/per L cell culture lysis buffer (1xPBS buffer (pH 7.4), 25% (w/v) sucrose, 5 mM ethylenediaminetetraacetic acid (EDTA), 1mM phenylmethylsulfonyl fluoride (PMSF)). Inclusion bodies were harvested by sonicating the cells on ice (5×30 s with 30 s rest between rounds at 80% amplitude), then centrifugation (25 min, 20000*g*, 4 °C) and the supernatant discarded. Resuspension, sonication, centrifugation, and discard of the supernatant was repeated twice, followed by resuspension of the pellet in 50 mM Tris-HCl (pH 7.4) and centrifugation (20 min, 20000*g*, 4 °C).

The pellet was resuspended in 12 mL 1.5% (w/v) sarkosyl, 10 mM dithiothreitol (DTT) and 20 mM Tris-HCl (pH 7.4) and incubated 3h at room temperature with mild agitation, followed by centrifugation (20 min, 20000*g*, 4 °C). The supernatant containing GST-SH was dialyzed against 0.5% (w/v) sarkosyl, 10 mM NaCl and 50 mM Tris-HCl (pH 7.4) at 4 °C to remove DTT and cleaved with thrombin (SERVA) to release GST, leaving two residues (GS) at the N terminus. Following cleavage, the solution was lyophilized and resuspended in 2 mL milliQ water. The solution was split into 100 μL aliquots to which each was added to 750 μL 1:2 chloroform:methanol solution and mixed well before addition of 300 μL milliQ water, the solution was mixed further and centrifuged (14000*g*, 2 min, 4 °C). Centrifugation resulted in three layers, the top aqueous layer was carefully removed and 500 μL methanol added to the solution, followed by mixing. The solution was incubated on ice for 20 min, centrifuged (16000*g*, 40 min, 4 °C), and the supernatant containing the target protein was transferred to a glass vial where the organic solvent was evaporated with a stream of N_2_.

The peptide SH_39-57_, Ac-RHAALYQRSFFHWSFDHSL, was acquired from TAG Copenhagen (DK) at 97% purity.

#### Peptide synthesis

The peptide SH_2-12_, H-Pro-Ala-Ile-Gln-Pro-Pro-Leu-Tyr-Leu-Thr-Phe-NH_2_, was synthesized using the Fmoc/t-Bu protection strategy on a Rink amide linker functionalized solid support, yielding the C-terminal amidated peptide after cleavage from the resin. Amino acids with acid-labile sidechain protecting groups were used, allowing simultaneous removal of protecting groups and release of peptide from the resin. The synthesis was conducted on an Initiator+ AlstraTM Automated Microwave Peptide Synthesizer (Biotage®). Each peptide coupling was performed twice at 75 °C for 5 minutes using diisopropylcarbodiimide (DIC)/Oxyma as the coupling reagent (5.00 eq./coupling), while Fmoc-deprotection was executed with 20% piperidine in DMF for 10 minutes at RT. Swelling and washing of the resin were carried out using dimethylmethanamide (DMF) at 70 °C for 20 minutes. Upon completion of the automated peptide synthesis, the resin was washed successively with DMF, MeOH, and CH_2_Cl_2_ before left with suction for 1 hour. Sidechain deprotection and cleavage of the peptide from the resin were achieved by adding trifluoroacetic acid (TFA)/H_2_O/ triisopropyl silane (TIPS) in a 95:2.5:2.5 ratio to the resin, followed by overnight shaking. The resulting suspension was filtered, and the filtrate was added cooled diethyl ether and TFA (TFA:diethyl ether 1:9). The precipitate was isolated by centrifugation and washed with diethyl ether (×3), then air-dried overnight. The crude peptide was divided into portions of maximum 25 mg, dissolved in a minimum volume of DMF, and purified by preparative HPLC. The pure fractions were collected, concentrated by air, and lyophilized, yielding the final pure peptide product. UPLC-MS *m/z*: 1274.2 [M+H^+^], calculated C_63_H_97_N_14_O_14_^+^: 1273.7. Automated solid-phase peptide synthesis was carried out using ChemMatrix resin beads as the solid support, with a loading of 0.3 mmol/g. The synthesis was performed in a flat-bottomed PE-syringe fitted with PPE filter, Teflon tubing and valve. Standard proteinogenic amino acids and the Rink amide linker were used as received from commercial sources. Preparative HPLC purification of the peptide was performed on a Waters auto-purification system consisting of a 2767 Sample Manager, 2545 Gradient Pump and 2998 PDA detector. Column: XBridge Peptide BEH C18 OBD Prep Column, 130 Å, 5 μm, 19 mm X 100 mm. Solvent A1: 0.1% formic acid in water; solvent B1: 0.1% formic acid in acetonitrile. Flow Rate: 20 mL/min.

#### Materials

DHPC, DPC, LMPG, 1-palmitoyl-2-hydroxy-sn-glycero-3-phospho-(1′-rac-glycerol) (LPPG), POPC and POPS were purchased from Avanti Polar Lipids. Sodium dodecyl sulphate (SDS) was purchased from Avantor Performance Materials. N-lauroylsarcosine sodium salt (sarkosyl), chloroform with 0.5-1.0 % ethanol as stabilizer, methanol ≥99.9 % and 2,2,2-Trifluoroethanol ≥99.0 % was purchased from Sigma-Aldrich.

#### Mass Spectrometry

SH_FL_ in DPC was buffer exchanged to 100 mM ammonium acetate and SH_1-34_ in DPC was buffer exchanged to 200 mM ammonium acetate with 2xCMC (0.046%) LDAO using Zeba biospin columns (7 kDa). The sample were further diluted to approximately 30 μM before it was transferred to a nESI capillary (Thermo). Using an off-line nanospray source, the sample was analysed on a Waters Synapt G1 TWIMS MS modified for analysis of intact protein complexes (MS Vision, The Netherlands) in positive ionization mode. The capillary voltage was 1.5 kV, the trap voltage 10V, and the cone voltage 10 V. The source temperature was 30°C. The source pressure was set to 8 mbar.

#### CD spectroscopy

CD spectra in the far (190–260 nm) UV ranges were recorded at 25 °C, 10 nm/min scan speed, 1 nm bandwidth and 2 s response using a Jasco J-810 or J-815 spectropolarimeter equipped with a sample holder Peltier temperature control and a path length of 0.1 cm using a quartz SUPRASIL cell (Hellma). For each condition, ten scans were accumulated, averaged and background spectrum recorded at identical conditions and subtracted. The spectra were smoothed with a convolution width of five data points. Concentrations of proteins and peptides varied between 5-30 μM in 20 mM Na_2_HPO_4_/NaH_2_PO_4_ (pH 7.2, or pH 6.5 for SH_34-57_), with detergents and lipids as indicated in the figure or figure legend. Mean residue ellipticity ([Ɵ_MRE_) was calculated using the following equation, where l is the path length in cm, c is the molar concentration and n is the number of peptide bonds (45).

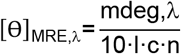

Fraction helicity was calculated based on the MRE at 222 nm, using the following equation:

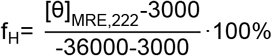

#### NMR spectroscopy

NMR samples for the detergent screen typically contained 200 μM SH_FL_ in 20 mM Na_2_HPO_4_/NaH_2_PO_4_ (pH 7.2) with 10% D_2_O (v/v) in the presence of a molar excess of detergents or lipids at an indicated ratio. Spectra for the assignment of backbone and sidechain nuclei were recorded on ^13^C,^15^N-labelled samples of 0.6 mM SH_FL_ in 66%TFE:23%H_2_O:10%D_2_O (v/v/v) and 0.8 mM SH_1-34_ in 1:500 DHPC in 20 mM Na_2_HPO_4_/NaH_2_PO_4_ (pH 7.2), 10% D_2_O (v/v) or on unlabelled samples of 0.6 mM SH_2-12_ in 20 mM Na_2_HPO_4_/NaH_2_PO_4_ (pH 7.2), 10% D_2_O (v/v) and 1 mM SH_39-57_ in milliQ, 10% D_2_O (v/v).

All NMR samples were recorded in 5 mm Shigemi BMS tubes (Bruker) and all NMR spectra were recorded on Bruker Avance III HD 600-, 750-, or 800-MHz spectrometers equipped with cryogenic probes and Z-field gradients at 37°C, except for SH_2-12_ recorded at 10°C. Free induction decays were transformed and visualized in NMRPipe (46) or Topspin (Bruker Biospin), and subsequently analysed using the CCPNmr Analysis software (47). Proton chemical shifts were internally referenced to DSS at 0.00 ppm, with heteronuclei referenced by relative gyromagnetic ratios. A ^1^H-^15^N HSQC spectrum was recorded for all samples. Further, for assignment of backbone and sidechain nuclei of ^13^C,^15^N-labbelled samples manual backbone assignment was performed based on the analysis of HNCO, HN(CA)CO, HNCA, HN(CO)CA, HNCACB, HN(CO)CACB spectra recorded with non-uniform sampling and processed using qMDD (48). For backbone assignment of unlabelled samples, natural abundance ^1^H-^1^H Total Correlated Spectroscopy (TOCSY) and ^1^H-^1^H Rotating frame Overhauser Effect Spectroscopy (ROESY) spectra were recorded, and manual backbone assignment performed. To acquire R1 and R2 relaxation rates for SH_1-34,_ triplicates of varying relaxation delays were recorded in a randomized order and peak intensities were fitted to single-exponential decay. Relaxation rates for T1 were 20 ms, 60 ms, 100 ms, 180 ms, 300 ms, 500 ms, 800 ms, 1200 ms, and for T2 16.96 ms, 32.92 ms, 67.84 ms, 101.76 ms, 135.68 ms, 169.6 ms, 203.52 ms, 271.36 ms. Dihedral angles for SH_1-34_ were calculated based on the chemical shift assignment using TALOS+ program (49).

Random coil chemical shifts for secondary structure analysis were generated using (https://www1.bio.ku.dk/english/research/bms/sbinlab/randomchemicalshifts2/) (50) and secondary chemical shift analysis was calculated using the following equation (51).

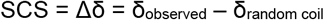

#### Model reconstruction

The pentameric assembly of the SH TM region was determined using the multibody docking capabilities of the HADDOCK webserver v.2.4 (23). A model of SH_7-34_ was generated using AlphaFold2. Five copies of SH_7-34_, where the AlphaFold model had a pLDDT score of ≥70, center-of-mass restraints and the dihedral angles obtained from NMR/TALOS imposed for the five monomers were used to obtain the TM pore. Center-of-mass restraints C5 symmetry restraints, and non-crystallographic restraints between the monomers were imposed. 10000 models were generated at the rigid body docking stage of which the best 400 were refined and clustered using the Fraction of Native Contacts (FCC) with a 0.6 cut-off. Clusters were ranked based on the default HADDOCK scoring function consisting of the intermolecular energies calculated using the OPLS force field (52) (100% van der Waals and 20% electrostatics) an empirical desolvation energy term (53) and the restraint energy. A 100 ns MD simulation of a peptide corresponding to the 23 C-terminal residues were performed using the CHARMM36 force field for GROMACS (54). Prominent conformations were split into eight clusters and three were removed due to steric hindrance for N-terminus of the peptide. Five copies of SH_35-57_ were added to multibody docking interface, restraining the distance of Lys35 from the peptides to Tyr34 of the TM helixes. Additionally, a shape consisting of fake beads was added to represent the phosphates in a POPC bilayer and Trp51 was ambiguously restrained to be within a 2 Å proximity to any shape bead. Peptide bonds were created between Lys35 in the C-terminal peptides and Tyr34 in the TM using PYMOL v. 2.5.5 and torsion angles were optimized in COOT. Similarly, the first 6 N-terminal residues were added from the AlphaFold3 model onto the pentameric model using PYMOL and torsion angles were optimized in COOT. A cross-section of the pentameric pore was analysed using the HOLE program (26).

#### Heterologous protein expression in *Xenopus l*. oocytes

Oocytes were purchased from Ecocyte (Germany) and kept in Kulori medium (in mm: 90 NaCl, 1 KCl, 1 MgCl_2_, 1 CaCl_2_, 5 HEPES adjusted to pH 7.4 with 2 m Tris-base ((HOCH_2_)_3_CNH_2_)) at 18°C. For oocyte experiments at least three preparations of oocytes from different frog donors were used. For the electrophysiological experiments, n is the number of oocytes. cDNA encoding C-terminally His6-tagged WT and truncated versions of SH Mumps (GenScript, Netherlands) in the pXOOM expression vector (55), linearized downstream from the poly-A segment, and *in vitro* transcribed using T7 mMessage machine in accordance with the manufacturer’s instructions (Ambion, Austin, TX, USA). cRNA was extracted with MEGAclear (Ambion) and stored at −80°C. For microinjection into defolliculated *Xenopus l*. oocytes, 50 ng of SH RNA/oocyte was used (4 ng RNA was also tested in the dose-dependent experiment), and the oocytes were kept for 3–4 days in Kulori medium at 18°C prior to experiments.

#### Two-electrode voltage clamp

Conventional two-electrode voltage clamp studies were performed using a DAGAN CA-1B High-Performance oocyte clamp (DAGAN, Minneapolis, MN, USA) with a Digidata 1440A interface controlled by pCLAMP software, version 10.5 (Molecular Devices, Burlingame, CA, USA). Electrodes were pulled (HEKA Elektronik, Lambrecht, Germany) from borosilicate glass capillaries to a resistance of 1.5–3 MΩ when filled with 1M KCl. The current traces were obtained by stepping the clamp potential from −20 mV to test potentials ranging from +50 mV to −130 mV (pulses of 200 ms) in increments of 15 mV. Recordings were low pass-filtered at 500 Hz, sampled at 1 kHz, and the steady-state current activity was analysed at 160–180 ms after applying the test pulse. The standard solution used consisted of (in mM) (100NaCl, 2KCl, 1CaCl_2_, 1MgCl_2_, 10 HEPES). Ion substitution experiments were measured on the same oocytes with or without the indicated ion, the Na^+^-free solution was made with equi-osmolar replacement with choline chloride, Ca^2+^-free solution was made with equi-osmolar replacement with Na^+^, and the Cl^-^-free solution was made with equi-osmolar replacement with Na^+^-gluconate. All solutions were adjusted to a pH of 7.4 with 2M Tris-base. To test inhibition by BIT225, oocytes expressing SH_FL_ were preincubated with BIT225 dissolved in DMSO (concentrations: 0.1, 3, 10 or 100 μM) with matched DMSO in control oocytes for 45 min prior to measurements.

#### Ligand synthesis

All reagents were of commercial grade. Silica gel 60 F254 precoated plates (Merck) were used for TLC. Flash chromatography of compounds was performed using silica gel 60 (40–64 μm). ^1^H NMR spectra were obtained at 400 or 600 MHz, and ^13^C NMR spectra were obtained at 100 or 151 MHz and calibrated relative to residual solvent peaks. Analytical HPLC was performed on a Dionex UltiMate system using a Gemini-NX C18 column (4.6 × 250 mm, 3 μm, 110 Å) with mobile phase A: water:TFA, 100:0.1, v/v, and mobile phase B: MeCN:water:TFA, 90:10:0.1, v/v/v. Data were acquired and processed using the Chromeleon Software v.6.80. The analysis was performed by a gradient of 0-100% mobile phase B in mobile phase A over 15 min.

Abbreviations: Acetyl (Ac), aqueous (aq.), broad (br), doublet (d), double doublet (dd), dichloromethane (DCM), dimethylformamide (DMF), dimethyl sulfoxide (DMSO), ethyl (Et), multiplet (m), methyl (Me), room temperature (rt), singlet (s), triplet (t), tetrahydrofuran (THF).

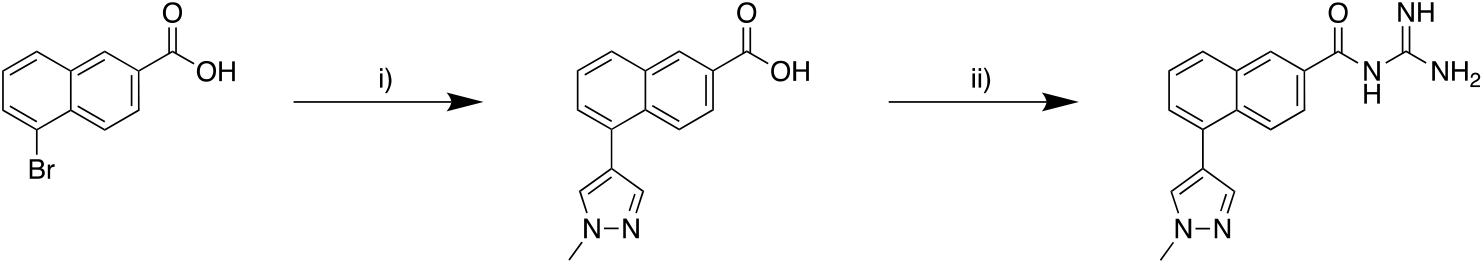

Reactions and Conditions: i) 1-methyl-1*H*-pyrazole-4-boronic acid pinacol ester, aq. NaOH (2.0 M), Na_2_CO_3_, MeCN, 90 °C, 18 h, 61%; ii) a) oxalyl chloride, DCM, DMF, rt (3.5 h)–40 °C (1 h), b) guanidine hydrochloride, aq. NaOH (2.0 M), THF, rt, 30 min, 65%.

5-(1-Methyl-1H-pyrazol-4-yl)-2-naphthoylguanidine (BIT225) (56) *Step 1:* 5-(1-Methyl-1H-pyrazol-4-yl)-2-naphthoic acid: A vial charged with 5-bromo-2-naphthoic acid (500 mg, 1.99 mmol), 1-methyl-1*H*-pyrazole-4-boronic acid pinacol ester (435.1 mg, 2.09 mmol), and Pd(PPh_3_)_4_ (115.1 mg, 99.6 μmol), was evacuated and backfilled with argon (3x). Then, aq. NaOH (2.0 M, 2.4 mL) and MeCN (9.5 mL) were added. The vial was sealed and stirred at 90 °C for 18 h. The reaction mixture was cooled down to rt and aq. HCl (1.0 M, 7.5 mL) was added, followed by water (5 mL) and extracted with EtOAc (3 × 15 mL). The combined organic phase was dried over anhydrous Na_2_SO_4_, filtered, and concentrated under reduced pressure. The crude product was recrystallised from DCM to afford a white solid (307 mg, 61%): *R*_f_ = 0.18 (MeOH:DCM); 5:95; ^1^H NMR (400 MHz, DMSO–*d*_6_) δ 13.14 (br s, 1H), 8.63 (d, J = 1.8 Hz, 1H), 8.22 (d, J = 8.9 Hz, 1H), 8.10 (s, 1H), 8.08 – 8.03 (m, 1H), 7.99 (dd, J = 8.8, 1.8 Hz, 1H), 7.75 (d, J = 0.8 Hz, 1H), 7.65 – 7.56 (m, 2H), 3.96 (s, 3H); ^13^C NMR (151 MHz, DMSO–*d*_6_) δ 167.4, 138.6, 132.8, 132.7, 131.0, 130.7, 130.3, 128.5, 128.3, 128.2, 126.5, 125.6, 125.5, 119.4, 38.7. *Step 2:* A flame-dried vial, charged with 5-(1-methyl-1H-pyrazol-4-yl)-2-naphthoic acid (100 mg, 0.40 mmol, was evacuated and backfilled with argon (3x). Then, anhydrous DCM (17 mL) and DMF (2 drops) were added followed by oxalyl chloride (100 μL). The mixture was stirred at rt for 3.5 h and then heated for 1 h at 40 °C. The cooled reaction mixture was concentrated under reduced pressure. The resulting crude acid chloride was suspended in anhydrous tetrahydrofuran (4 mL) and this mixture was added dropwise to a solution of guanidine hydrochloride (174.2 mg, 1.82 mmol) in aq. NaOH (2.0 M, 2.4 mL, 4.8 mmol) and the reaction mixture was then stirred for 30 min. Water (3 mL) was added and extracted with EtOAc (3 × 5 mL). The combined organic phase was washed sequentially with aq. NaOH (1 M, 10 mL) and water (10 mL), then dried over anhydrous Na_2_SO_4_ and filtered. The filtrate was evaporated onto Celite and purified by flash chromatography (3-10% EtOAc in heptane) to afford the title compound as a white solid (75 mg, 65%). *R*_f_ = 0.23 (MeOH:DCM; 1:9); ^1^H NMR (600 MHz, CD_3_OD) δ 8.65 – 8.60 (m, 1H), 8.13 (d, J = 1.2 Hz, 2H), 7.94 – 7.88 (m, 2H), 7.73 (s, 1H), 7.54 – 7.49 (m, 2H), 4.02 (s, 3H); ^13^C NMR (151 MHz, CD_3_OD) δ 179.4, 165.0, 140.1, 137.2, 134.8, 134.3, 131.8, 130.8, 129.6, 129.2, 127.0, 126.9, 126.0, 122.3, 39.0; HPLC: *t*_R_ = 9.12 min, purity: 96.3%.

## Statistical Analyss

Statistical analyses were performed with Prism7 (GraphPad Software Inc., La Jolla, CA, USA), and the test stated in the figure legends. An outlier was excluded from Fig. 4i, after Grubbs test. Data represent the mean ± SEM and P < 0.05 was considered statistically significant.

## Data availability

NMR chemical shifts have been deposited in BioMagResBank under the accession numbers 52513, 52514, 52515 and 52516. HADDOCK json files and PDB files of the structural models have been deposited to Zenodo (DOI 10.5281/zenodo.12698126).

