## Supplementary figures for "The SH Protein of Mumps Virus is a Druggable Pentameric Viroporin"

Devantier et al., 2024

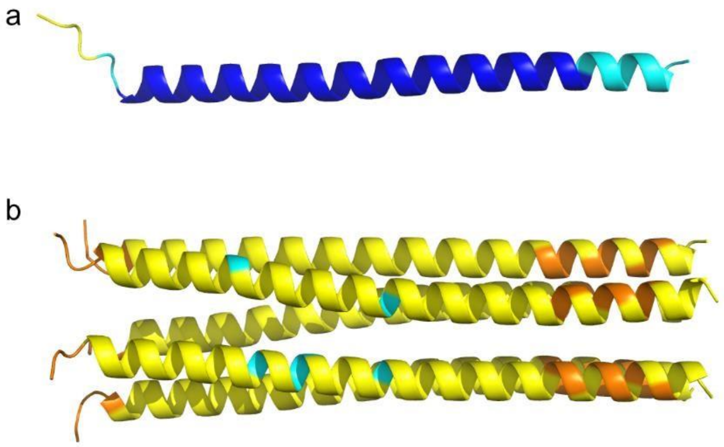


**Fig. S1. AlphaFold3 models of $SH_{FL}$.** a) Monomer. b) Pentamer. Coloured according to the pLDDT score (yellow <70, blue 70-100).

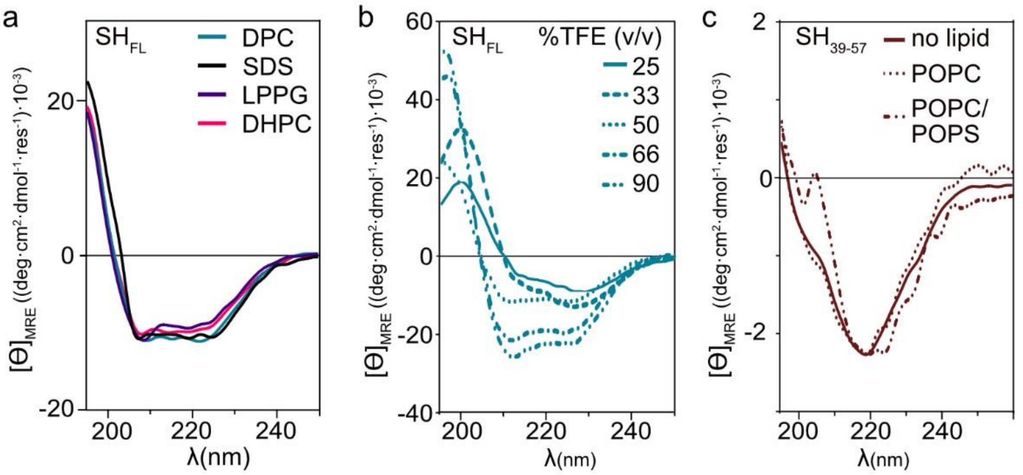


**Fig. S2. Effects of membrane mimetics on $SH_{FL}$ and $SH_{39-57}$.** a) Far-UV CD spectra of $SH_{FL}$ in detergents as indicated. b) FarUV CD spectra of induced helix in $SH_{FL}$ with added TFE content. c) Far-UV CD spectra of $SH_{39-57}$ with lipids as indicated, normalized to MRE at 218 for $SH_{39-57}$ with no lipids.

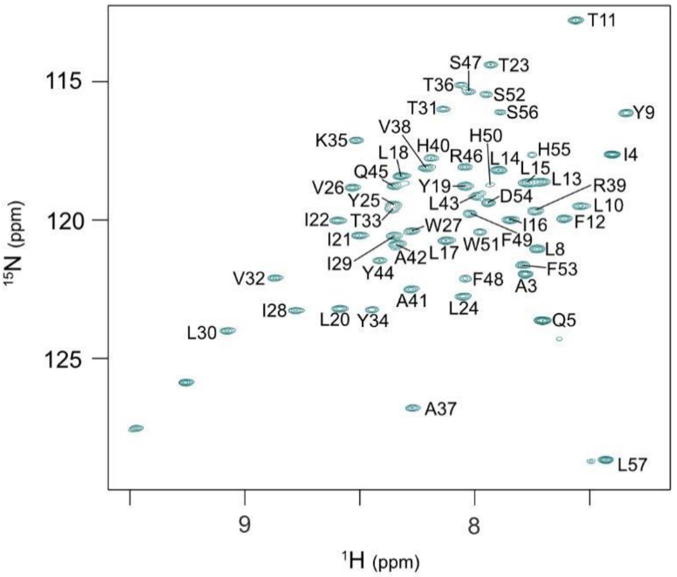


**Fig. S3. $SH_{FL}$ in 66% TFE:33%$H_2O$.** $^1$H-$^{15}$N HSQC spectrum of $SH_{FL}$ in 66% TFE:33% $H_2O$, with assignment.

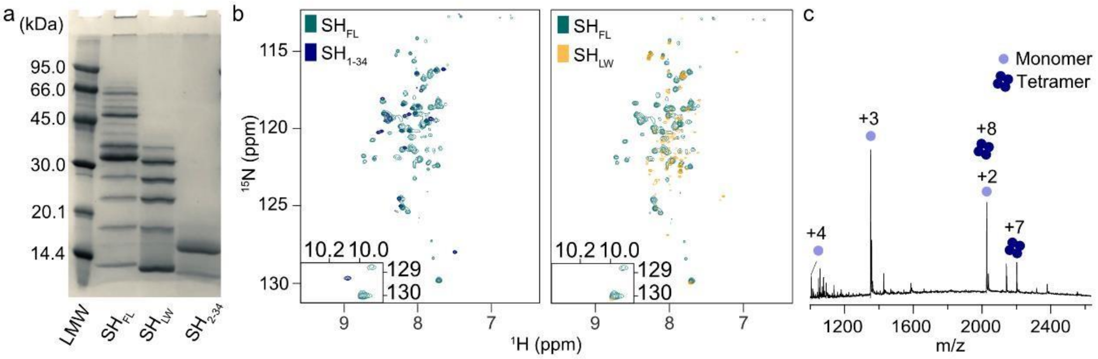


**Fig. S4. SH variants.** a) SDS-PAGE of SH variants in 100 mM SDS. b) Left: $^1$H-$^{15}$N HSQC spectra of $SH_{FL}$ (green) and $SH_{1\text{-}34}$ (blue) in 1:500 DHPC. Right: $^1$H-$^{15}$N HSQC spectra of $SH_{FL}$ (green) and $SH_{LW}$ (yellow) in 1:500 DHPC. c) Native mass spectrometry of $SH_{1\text{-}34}$.

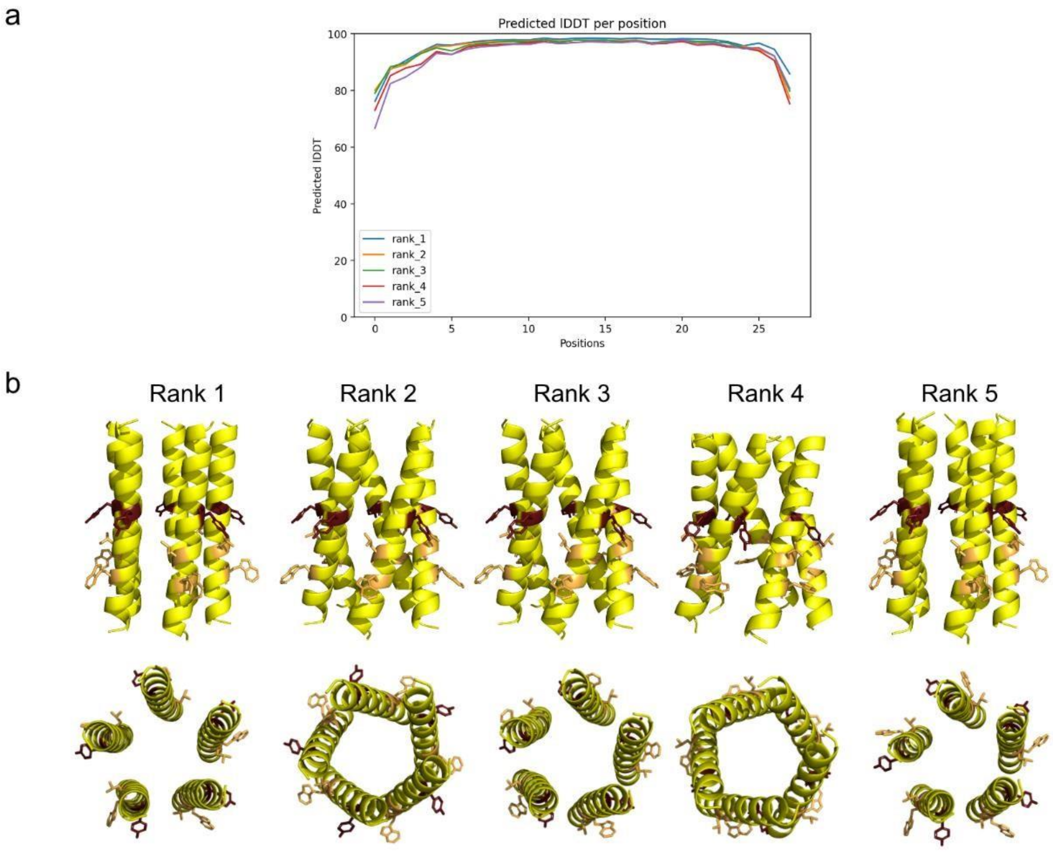


**Fig. S5. $SH_{7\text{-}34}$ AlphaFold models**. a) pLDDT scores for the five predicted AlphaFold2 monomeric models of residues $SH_{7\text{-}34}$, labelled position 1-27. b) The five pentameric models generated by AlphaFold3. Above: Side view. Below: N-terminal view. Coloured according to the pLDDT score (yellow <70, blue 70-100). Residues Tyr19 (red), Leu24 (yellow), and Trp27 (yellow) shown as sticks.

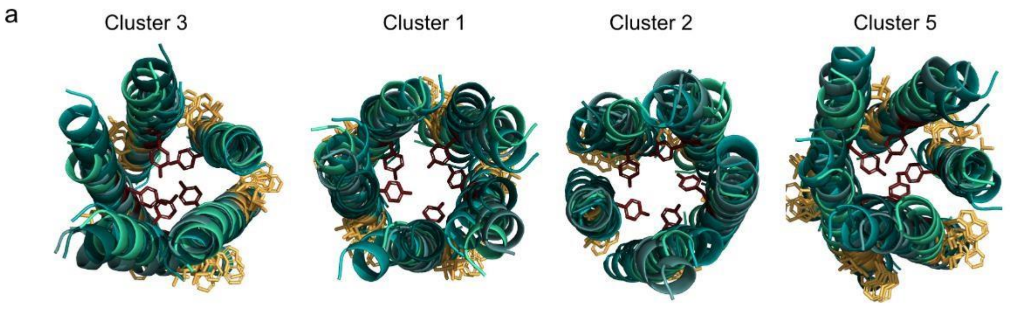


b

| | HADDOCK Score | RMSD-$E_{min}$ | N | $E_{vdw}$ | $E_{elec}$ | BSA (Å$^2$) | $E_{desolv}$ |
|---|---|---|---|---|---|---|---|
| 3 | -418 (14) | 13.8 (0.2) | 7 | -178 (5) | -39 (11) | 6462 (174) | -232 (8) |
| 1 | -412 (18) | 5.5 (1.4) | 10 | -170 (10) | -31 (21) | 6274 (249) | -236 (12) |
| 2 | -402 (32) | 2.3 (1.3) | 8 | -166 (24) | -21 (9) | 6050 (301) | -233 (11) |
| 5 | -390 (33) | 14.1 (0.1) | 5 | -165 (20) | -31 (8) | 5954 (268) | -219 (15) |

**Fig. S6. HADDOCK output for $SH_{7\text{-}34}$**. a) N-terminal view of top 4 clusters. Residues Tyr19 (red), Leu24 (yellow), and Trp27 (yellow) shown as sticks. b) Table of Haddock statistics with standard deviations indicated between brackets. The HADDOCK score and various other reported statistics are calculated as the average over the top 4 members of a cluster. N indicates the number of models in each cluster. $E_{vdw}$, $E_{elec}$, BSA, and $E_{desolv}$ represent the intermolecular van der Waals and electrostatic energies, the buried surface area, and the desolvation energy, respectively.

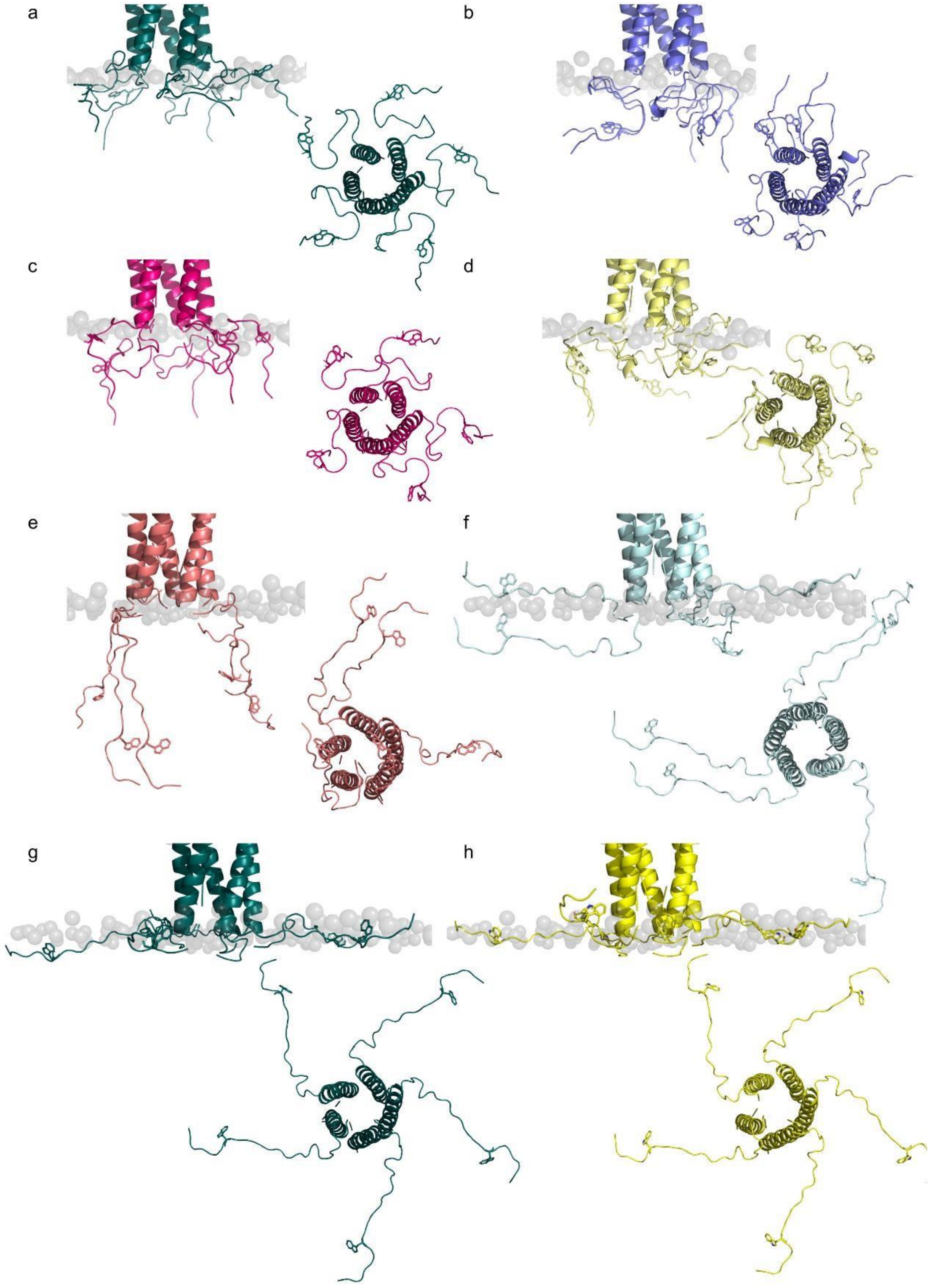


**Fig. S7. Representative HADDOCK output for the SH-pentamer and C-terminal peptide docking.** Side and C-terminal view of top clusters ordered by ranking (a-h). Residues Trp51 is shown as sticks.
